# Polygenic scoring accuracy varies across the genetic ancestry continuum in all human populations

**DOI:** 10.1101/2022.09.28.509988

**Authors:** Yi Ding, Kangcheng Hou, Ziqi Xu, Aditya Pimplaskar, Ella Petter, Kristin Boulier, Florian Privé, Bjarni J. Vilhjálmsson, Loes Olde Loohuis, Bogdan Pasaniuc

**Affiliations:** Bioinformatics Interdepartmental Program, UCLA, Los Angeles, CA, USA; Department of Computer Science, UCLA, Los Angeles, CA, USA; National Center for Register-Based Research, Aarhus University, Aarhus, Denmark; Bioinformatics Research Centre, Aarhus University, Aarhus, Denmark; Novo Nordisk Foundation Center for Genomic Mechanisms of Disease, Broad Institute, Cambridge, MA, USA; Department of Psychiatry, David Geffen School of Medicine at UCLA, Los Angeles, CA, USA; Program in Neurobehavioral Genetics, David Geffen School of Medicine at UCLA, Los Angeles, CA, USA; Department of Computational Medicine, David Geffen School of Medicine at UCLA, Los Angeles, CA, USA; Department of Human Genetics, David Geffen School of Medicine at UCLA, Los Angeles, CA, USA; Department of Pathology and Laboratory Medicine, David Geffen School of Medicine at UCLA, Los Angeles, CA, USA

## Abstract

Polygenic scores (PGS) have limited portability across different groupings of individuals (e.g., by genetic ancestries and/or social determinants of health), preventing their equitable use. PGS portability has typically been assessed using a single aggregate population-level statistic (e.g., R^2^), ignoring inter-individual variation within the population. Here we evaluate PGS accuracy at individual-level resolution, independent of its annotated genetic ancestries. We show that PGS accuracy varies between individuals across the genetic ancestry continuum in all ancestries, even within traditionally “homogeneous” genetic ancestry clusters. Using a large and diverse Los Angeles biobank (ATLAS, N= 36,778) along with the UK Biobank (UKBB, N= 487,409), we show that PGS accuracy decreases along a continuum of genetic ancestries in all considered populations and the trend is well-captured by a continuous measure of genetic distance (GD) from the PGS training data; Pearson correlation of −0.95 between GD and PGS accuracy averaged across 84 traits. When applying PGS models trained in UKBB “white British” individuals to European-ancestry individuals of ATLAS, individuals in the highest GD decile have 14% lower accuracy relative to the lowest decile; notably the lowest GD decile of Hispanic/Latino American ancestry individuals showed similar PGS performance as the highest GD decile of European ancestry ATLAS individuals. GD is significantly correlated with PGS estimates themselves for 82 out of 84 traits, further emphasizing the importance of incorporating the continuum of genetic ancestry in PGS interpretation. Our results highlight the need for moving away from discrete genetic ancestry clusters towards the continuum of genetic ancestries when considering PGS and their applications.

## Introduction

Polygenic scores (PGS)—estimates of an individual’s genetic predisposition for complex traits/diseases (i.e. genetic value)—are a promising application of large-scale genome-wide association studies (GWAS) to personalized genomic medicine^1–4^, disease risk prediction and prevention^5–8^. The portability of PGS across different ancestry and socio-demographic groups is limited due to Euro-centric sampling of GWAS data coupled with differences in linkage disequilibrium (LD), minor allele frequency (MAF) and/or disease genetic architecture^3,9–13^, which poses a critical equity barrier that has prevented widespread adoption of PGS for personalized medicine. For example, PGS are significantly more accurate for individuals of European ancestries as compared to other genetic ancestries^10,14^; furthermore, PGS accuracy varies across socio-genomic features (e.g., sex, age and social economic status)^11^, thus complicating interpretability of PGS across groups with different environmental exposures.

PGS accuracy is traditionally assayed using population-level metrics of accuracy (e.g., R^2^), thus assuming some level of homogeneity across individuals within the considered population^2,11,15^. However, homogeneous populations are an idealized concept that only roughly approximate human populations; human diversity exists along a genetic ancestry continuum without clearly defined clusters and with various correlations between genetic and socio-environmental factors^15–20^. Grouping individuals into genetic ancestry clusters obscures the impact of individual variation on PGS accuracy. This is evident for individuals with recently admixed genomes, where genetic ancestries vary individual-to-individual and locus-to-locus in the genome. For example, a single population-level PGS accuracy estimated across all African Americans greatly overestimates PGS accuracy for African Americans with large proportions of African genetic ancestries^21^; likewise, coronary artery disease PGS performs poorly in Hispanic individuals with high proportions of African ancestry^22^. The genetic ancestry continuum impacts PGS accuracy even in traditionally-labeled “homogeneous/non-admixed” populations; for example, PGS accuracy decays across a gradient of subcontinental ancestries within Europe as the target cohorts are more genetically dissimilar from the data used to train the PGS^19,23^. Assessing PGS accuracy using population-level metrics is further complicated by technical issues in assigning individuals to discrete clusters of genetic ancestries. Different algorithms and/or reference panels may assign the same individual to different clusters^15,23,24^ and thus to different PGS accuracy classes. Moreover, many individuals are not assigned to a cluster due to limited reference panels used for genetic ancestry inference^23,25^, leaving such individuals outside PGS accuracy characterization; this poses equity concerns as it limits PGS applications only to individuals within well-defined clusters of genetic ancestries.

Here we leverage methods that characterize PGS performance at the level of a single target individual^26^ to evaluate the impact of the genetic ancestry continuum on PGS accuracy. We use simulation and real data analysis to show that PGS accuracy decays continuously individual-to-individual across the genetic continuum of ancestry as function of genetic distance (GD) from the PGS training data; GD is defined as a principal component analysis (PCA) projection of the target individual on the training data used to estimate the PGS weights. We leverage a large and diverse Los Angeles biobank at UCLA (ATLAS, N= 36,778) joint with UK Biobank (UKBB, N= 487,409) to investigate the interplay between genetic ancestries and PGS for 84 complex traits and diseases. The accuracy of PGS models trained in UKBB “white British” individuals (N= 371,018) is negatively correlated with GD for all considered traits (average Pearson R= −0.95 across 84 traits), demonstrating pervasive individual variation in PGS accuracy. The negative correlation remained significant even when restricted to traditionally defined “homogenous” clusters of genetic ancestries (ranging from R=-0.43 for East Asian cluster to R=-0.85 for the African American cluster in ATLAS). On average across the 84 traits, when rank-ordering individuals according to distance from training data, PGS accuracy decreased by 14% in the closest vs furthest decile in the European genetic cluster; notably the furthest decile of European ancestry individuals showed similar accuracy to the closest decile of Hispanic Latino individuals. Characterizing PGS accuracy across continuum of GD allows for inclusion of individuals unassigned to a given genetic ancestry clusters (6% of all ATLAS), thus allowing for more individuals to be included in PGS applications. Finally, we explore the relationship between GD and PGS estimates themselves; 82 (out of 84) PGS show significant correlation between GD and PGS with 30 showing opposite correlation (GD, trait) vs (GD, PGS); we exemplify the importance of incorporating GD in interpretation of PGS using height and neutrophils in the ATLAS data. Overall, our results demonstrate the need to incorporate the genetic ancestry continuum on PGS performance and/or bias.

## Results

### Overview of the study

PGS accuracy has traditionally been assessed at the level of discrete genetic ancestry clusters using population-level metrics of accuracy (e.g., R^2^). Individuals from diverse genetic backgrounds are routinely grouped into discrete genetic ancestry clusters using computational inference methods such as PCA^27^ and/or admixture analysis^28^ (Figure 1a). Population-level metrics of PGS accuracy are then estimated for each genetic ancestry cluster and generalized to each individual in the cluster (Figure 1b). This approach has three major limitations: (1) the inter-individual variability within each cluster is ignored; (2) the genetic ancestry cluster boundary is sensitive to algorithms and reference panels used for clustering; and (3) a significant proportion of individuals may not be assigned to any cluster due to a lack of reference panels for genetic ancestry inference (e.g., individuals of uncommon or admixed ancestries).

**Figure 1.**
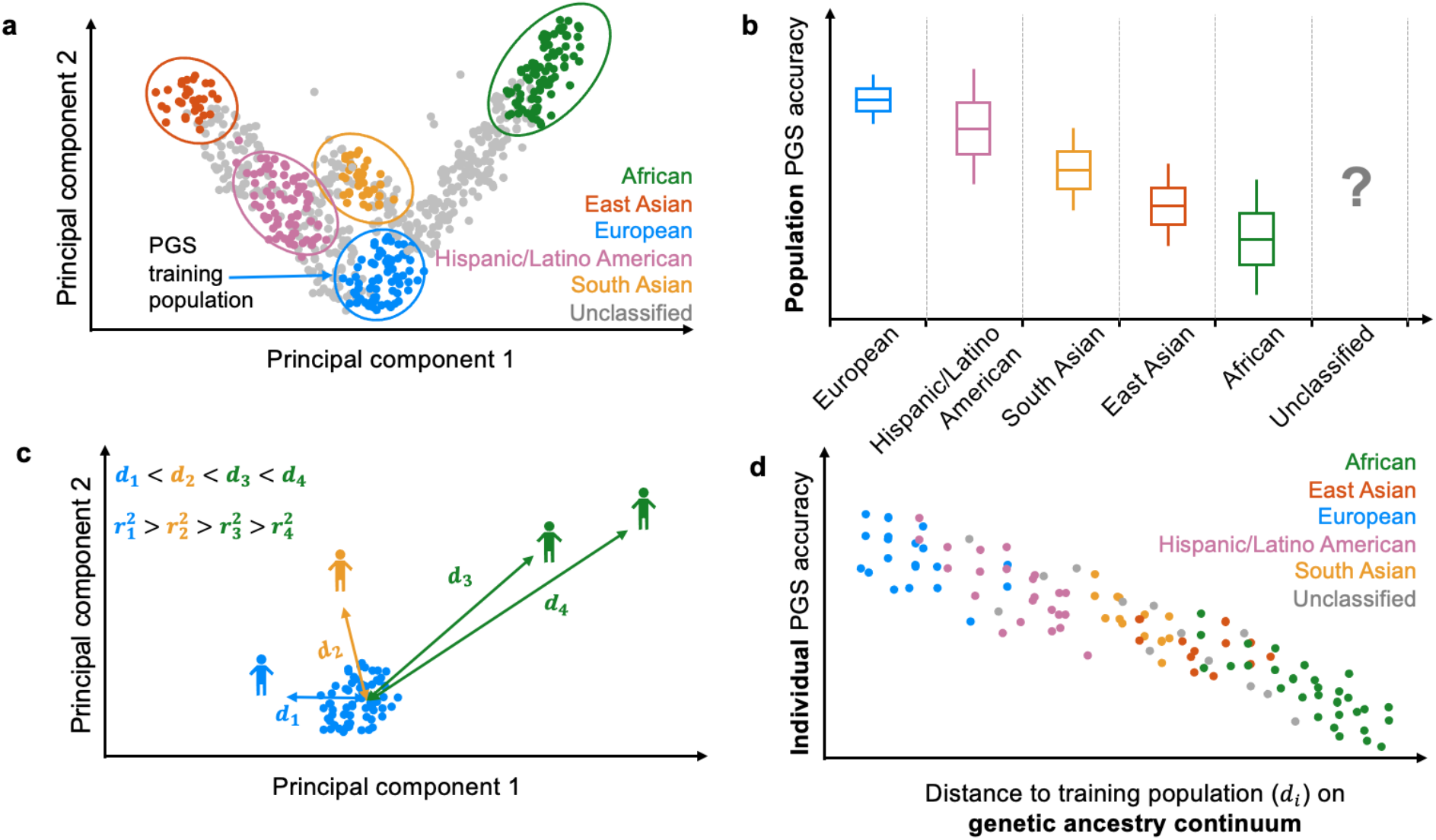
Population-level vs individual-level PGS accuracy. (a) Discrete labeling of genetic ancestry with PCA-based clustering. Each dot represents an individual. The circles represent arbitrary boundaries imposed on the genetic ancestry continuum to divide individuals into different genetic ancestry clusters. The color represents the assigned genetic ancestry cluster label. The gray dots are individuals who are left unclassified. (b) Population-level PGS accuracy varies across clusters. The box plot represents the PGS accuracy (e.g., R2) measured at population level. The question mark emphasizes that the PGS accuracy for unclassified individuals is unknown due to the lack of a reference group. Gray dashed lines emphasize the categorical nature of genetic ancestry clustering. (c) Continuous labeling of each individual’s unique position on genetic ancestry continuum with a PCA-based genetic distance. The genetic distance is defined as the Euclidean distance of an individual’s genotype from the center of the training data when projected on the PC space of training genotype data. Each individual has its own unique genetic distance *d_i_* and individual PGS accuracy 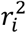. (d) Individual-level PGS accuracy decays along the genetic ancestry continuum. Each dot represents an individual and its color represents the genetic ancestry label assigned in panel a. Individuals labeled with the same ancestry spread out on the genetic ancestry continuum and there are no clear boundaries between genetic ancestry groups.

In this work, we evaluate PGS accuracy across the genetic ancestry continuum at level of a single target individual. We model the phenotype of individual *i*. as 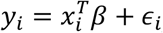, where *x_i_* is a *M* × 1 genotype vector indicating allele counts, *β* is a *M* × 1 allelic causal effects vector and *ϵ_i_* is random noise. Under a random effects model 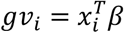 and 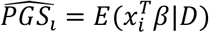 are random variables where the randomness comes from *β* and training data *D* (*D* = (*X_train_*, *y_train_*)). We define the individual PGS accuracy as the correlation of an individual’s genetic value and PGS estimates as:

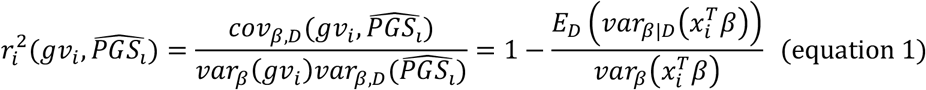

We use Ldpred2 to estimate 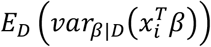^26,29^ and approximate 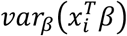 as the heritability of the phenotype (Methods)^30^; equation 1 can be further simplified assuming all variants are causal drawn from a normal distribution (infinitesimal model, see Methods). As continuous genetic distance (GD) we use 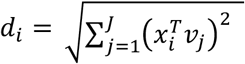 where *v_j_* is the *j_th_* eigenvector of training genotype data (Figure 1c). Individuals that are clustered into the same genetic ancestry clusters may have different genetic distance from training data and different individual PGS accuracy (Figure 1d). We use theory and empirical data analyses to show that PGS accuracy decay is well-approximated by the continuous metric of genetic distance.

We organize the manuscript as follows. First, we show the relation between genetic distance and PGS accuracy in simulations using real genotype data from UK biobank. Next, we show that existing PGS have accuracy that decreases individual-to-individual as function of genetic distance in a diverse biobank from UCLA. Finally we showcase the impact of genetic distance on interpretability of PGS using height and neutrophil count as example traits.

### Individual PGS performance is calibrated across the genetic ancestry continuum in simulations

First, we evaluated calibration of 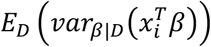 estimated by LDpred2 for individuals at various genetic distances from the UKBB “white British” individuals used to train PGS by checking the calibration the of 90% credible intervals (Figure 2a). We simulated 100 phenotypes at heritability 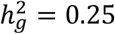 and proportion of causal variants *p_causal_* = 1% for all individuals in UK Biobank, assuming shared causal variants and homogenous allelic effect sizes for individuals from various genetic backgrounds (see Methods). Overall, the 90% credible intervals are approximately well-calibrated, i.e. the credible interval overlaps with the true genetic value across 90 out of 100 replicates, for all individuals, regardless of their distance from the training population or genetic ancestry labels (Figure 2a). For example, when individuals are binned into 10 deciles based on their GD from the training population, the average empirical coverage of the 90% credible intervals is 89.7% (s.d. 2.6%) for individuals from the lowest decile (composed of 96.9% individuals labeled as “white British”, 3.1% labeled as “Poland” under discrete view of ancestries) compared with the average empirical coverage of 82.4 % (s.d. 4.6%) for individuals from the highest decile (composed of 19.9% individuals labeled as “Caribbean” and 80.1% labeled as “Nigeria”).

**Figure 2.**
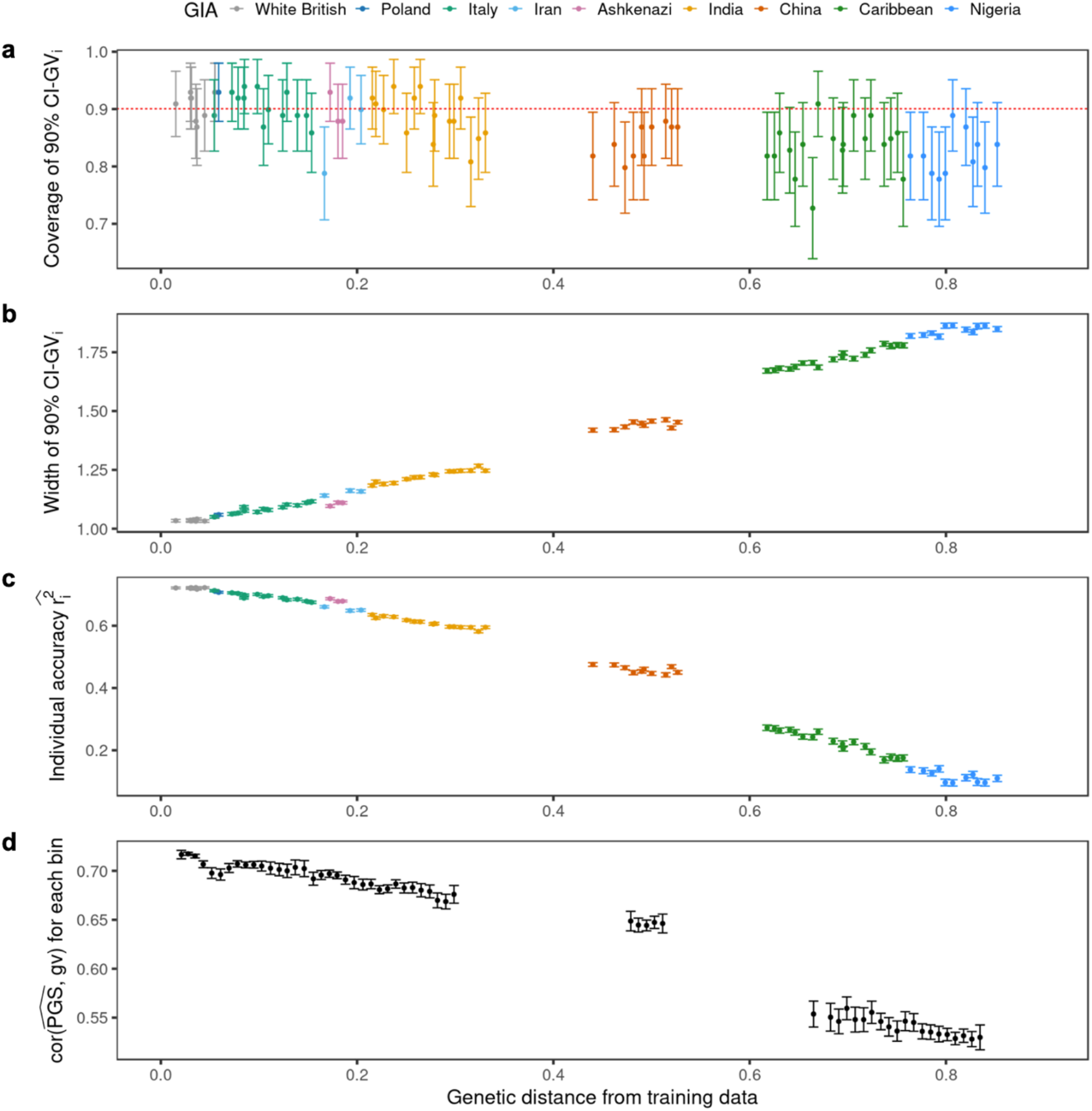
PGS performance is calibrated across genetic distance in simulations using UKBB data. (a) 90% credible intervals are well calibrated for testing individuals at all genetic distances. The red dotted line represents the expected coverage of 90% credible interval. Each dot represents a randomly selected UKBB testing individual. For each dot, the x-axis is its genetic distance from training data, the y-axis is the empirical coverage of 90% credible interval calculated as the proportion of simulation replicates where the 90% credible intervals contain the individual’s true genetic value, and the error bars represent mean ±1.96 standard error of the mean (s.e.m) of the empirical coverage calculated from 100 simulations. (b) The width of 90% credible interval increases with genetic distance. For each dot, the y-axis is the width of 90% credible interval across 100 simulation replicates, and the error bars represent ±1.96 s.e.m. (c) Individual PGS accuracy decreases with genetic distance. For each dot, the y-axis is the average individual level PGS accuracy across 100 simulation replicates, and the error bars represent ±1.96 s.e.m. (d) Population-level metrics of PGS accuracy recapitulates the decay in PGS accuracy across genetic continuum. All UKBB testing individuals are divided into 100 equal-interval bins based on their genetic distance. The x-axis is the average genetic distance for the bin and the y-axis is the squared correlation between genetic value and PGS estimates for the individuals within the bin. The dot and error bars represent the mean and ±1.96 s.e.m from 100 simulations.

Next, we investigated the impact of GD on individual-level PGS accuracy. As expected, the credible interval width increases linearly with GD reflecting reduced predictive accuracy for the PGS (Figure 2b). The average width of 90% credible interval is 1.83 in the highest decile of GD, a 1.8-fold increase over the average width in the lowest decile of GD. In contrast to the credible interval width, the individual-level PGS accuracy 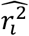 decreases with genetic distance from training data (Figure 2c); the average estimated accuracy of individuals in the lowest decile GD is 4-fold higher than that of individuals in the highest decile. Even among the most homogenous grouping of individuals traditionally labeled as white British, we observe a 5% relative decrease in accuracy for individuals at the highest decile of GD as compared to those in the lowest decile. Similar results are observed when using population-level PGS metric of accuracy, albeit at expense of binning individuals according to GD; we find a high degree of concordance between the average 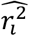 within the bin and the population-level R^2^ estimated within the bin (Figure 2d, Supplementary Figure 1a). Similarly, we observe a high consistency between average 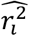 and squared correlation between PGS and simulated phenotype, (R = 0.87, P < 2.2e-16, Supplementary Figure 1b). Taken together, our results show that 90% credible interval remains calibrated for individuals that are genetically distant from the training population at the expense of a wider credible intervals while 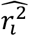 captures the PGS accuracy decay across genetic distance.

### Individual PGS accuracy varies across the genetic ancestry continuum in all ancestries

After having validated our approach in simulations, we next turn to empirical data. For illustration purposes we use height as example focusing on ATLAS biobank as target population with PGS trained in the 371,018 “white British” individuals from UKBB (Methods); other traits show similar trends and are presented in the next sections. PGS accuracy at the individual level varies with GD across the entire biobank as well as within all genetically inferred ancestry clusters (Figure 3, Supplementary Fig 2). For example, GD strongly correlates with PGS accuracy of individuals in the genetic ancestry cluster labeled as Hispanic/Latino American (HL, R = −0.83) and African American (AA, R=-0.88) in ATLAS. Notably, GD correlates with PGS accuracy even in non-admixed genetic clusters of ancestry with correlations as −0.66, −0.66 and −0.35, for European Americans (EA), South Asian Americans (SAA) or East Asian Americans (EAA), respectively. Similar qualitative results are also observed when applying PGS in a test data from UKBB with significant negative correlations between GD and individual PGS accuracy in all the sub-continental genetic clusters in UKBB (Supplementary Figure 2) ranging from R= −0.031 for the “white British” cluster to R=-0.62 for the Caribbean cluster.

**Figure 3.**
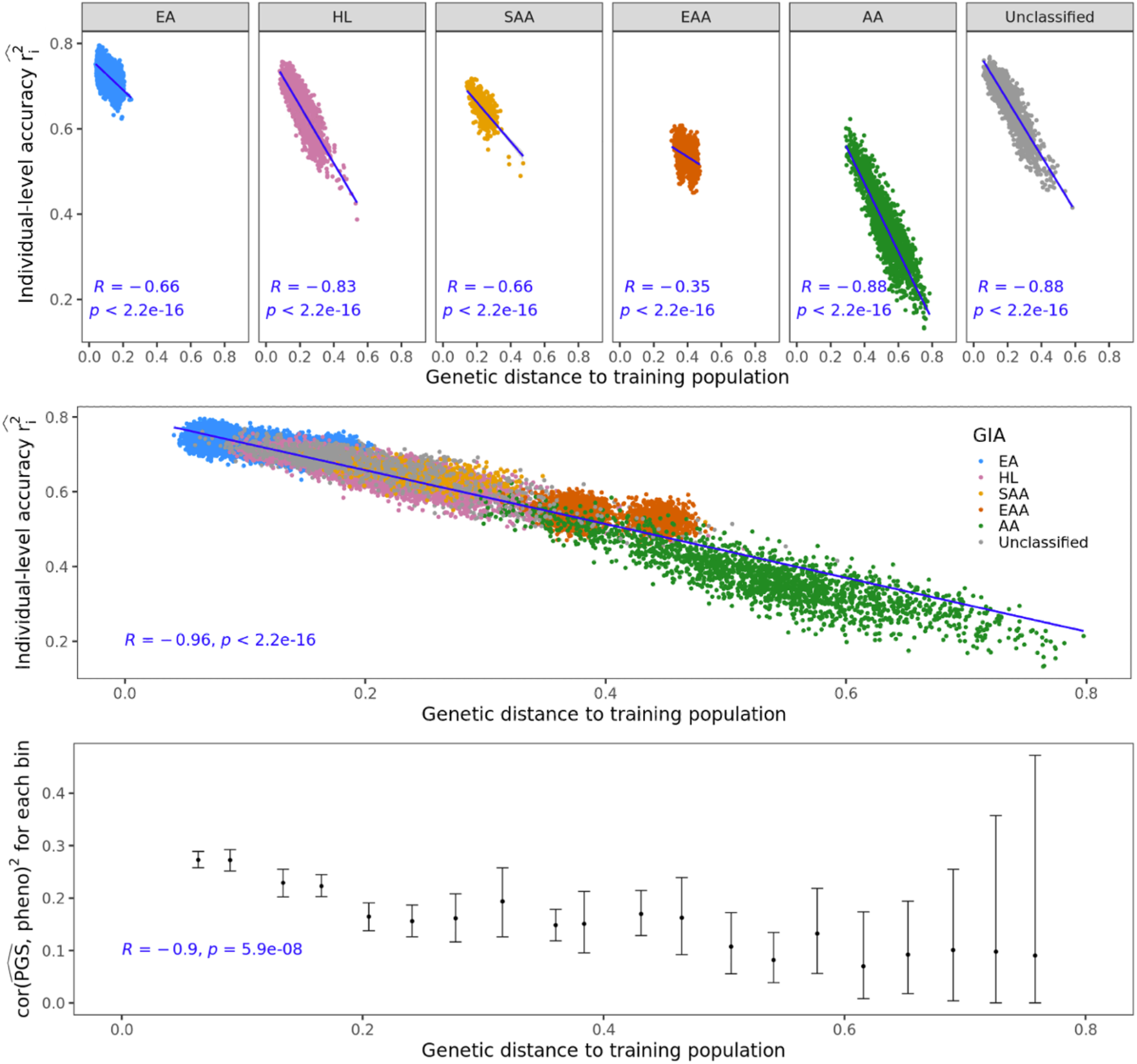
The individual-level accuracy for height PGS decreases across the genetic ancestry continuum in ATLAS. (a) Individual PGS accuracy decreases within both homogenous and admixed genetic ancestry clusters. Each dot represents a testing individual from ATLAS. For each dot, the x-axis represents its distance from the training population on the genetic continuum; the y-axis represents its PGS accuracy. The color represents the inferred genetic ancestry cluster. R and p refer to the correlation between genetic distance and individual-level PGS accuracy and its significance from two-sided t-tests. (b) Individual PGS accuracy decreases across the entire ATLAS. (c) Population-level PGS accuracy decreases with the average genetic distance in each genetic distance bin. All ATLAS individuals are divided into 20 equal-interval genetic distance bins. The x-axis is the average genetic distance within the bin, the y-axis is the squared correlation between PGS and phenotype for individuals in the bin; The dot and error bar show mean and 95% confidence interval from 1000 bootstrap samples. (EA, European American; HL, Hispanic/Latino American; SAA, South Asian American; EAA, East Asian American; AA, African American.)

Next, we focused on the impact of GD on PGS accuracy across all ATLAS individuals regardless of genetic ancestry clustering (R = −0.96, P < 2.2 e-16, Figure 3b). Notably, we find a strong overlap of PGS accuracies across individuals from different genetical ancestry clusters demonstrating the limitation of using a single cluster-specific metric of accuracy. For example, when rank-ordering by GD, we find the individuals from the closest GD decile in HL cluster have similar estimated accuracy as the individuals from the farthest GD decile in EA cluster (average 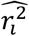 of 0.71 vs 0.71). This shows that GD enables identification of HL individuals with similar PGS performance as the EA cluster thus partly alleviating inequities due to lack of access to accurate PGS. Most notably, GD can be used to evaluate PGS performance for individuals that cannot be easily clustered by current genetic inference methods (6% of all individuals in ATLAS, Figure 3b) partly due to limitations of reference panels and algorithms for assigning ancestries. Among this traditionally overlooked group of individuals, we find the GD ranging from 0.02 to 0.64 and their corresponding estimate PGS accuracy 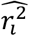 ranging from 0.63 to 0.21. In addition to evaluating PGS accuracy with respect to the genetic value, we also evaluated accuracy with respect to the residual height after regressing out sex, age, PC1-10 on the ATLAS from the actual measured trait. Using equally spaced bins across the GD continuum, we find that correlation between PGS and the measured height tracks significantly with GD (R = −0.9, P-value = 5.9e-8, Figure 3c).

### The continuous decay of PGS accuracy across genetic distance is pervasive across all traits

Having established the coupling of GD with PGS accuracy in simulations and for height, we next turn to the question of whether such relationship is pervasive across complex traits using PGS for a broad set of 84 traits (Supplementary Table 1). We find consistent and pervasive correlations of GD with PGS accuracy across all considered traits in both ATLAS and UK Biobank (Figure 4). For example, the correlations between GD and individual PGS accuracy range from −0.71 to −0.97 with an average of −0.95 across the 84 PGS in ATLAS with similar results in UKBB. Traits with sparser genetic architectures and fewer non-zero weights in the PGS yield to a lower correlation between GD and PGS accuracy; we hypothesize this is because GD represents genome-wide genetic variation patterns that may not reflect a limited number of causal SNPs well. For example, PGS for Lipoprotein A (log_lipoA) has the lowest polygenicity estimate (0.02%) among the 84 traits and has the lowest correlation in ATLAS (−0.71) and UKBB (−0.85). In contrast, we observe a high correlation between GD and PGS accuracy (>0.9) for all traits with an estimated polygenicity > 0.1%. Next, we show that the fine-scale population structures accountable for the individual PGS accuracy variation is also prevalent within the traditionally defined genetic ancestry group. For example, in ATLAS we find 501 out of 504 (84 traits across 6 genetic ancestry clusters) trait-ancestry pairs have a significant association between GD and individual PGS accuracy after Bonferroni correction. In UKBB, we find 572 out of the 756 (84 traits across 9 subcontinental genetic ancestry clusters) trait-ancestry pairs have significant association between genetic distance and PGS accuracy after Bonferroni correction. We also find that a more stringent definition of homogenous genetic clusters results in a lower correlation magnitude (Supplementary Figure 3).

**Figure 4.**
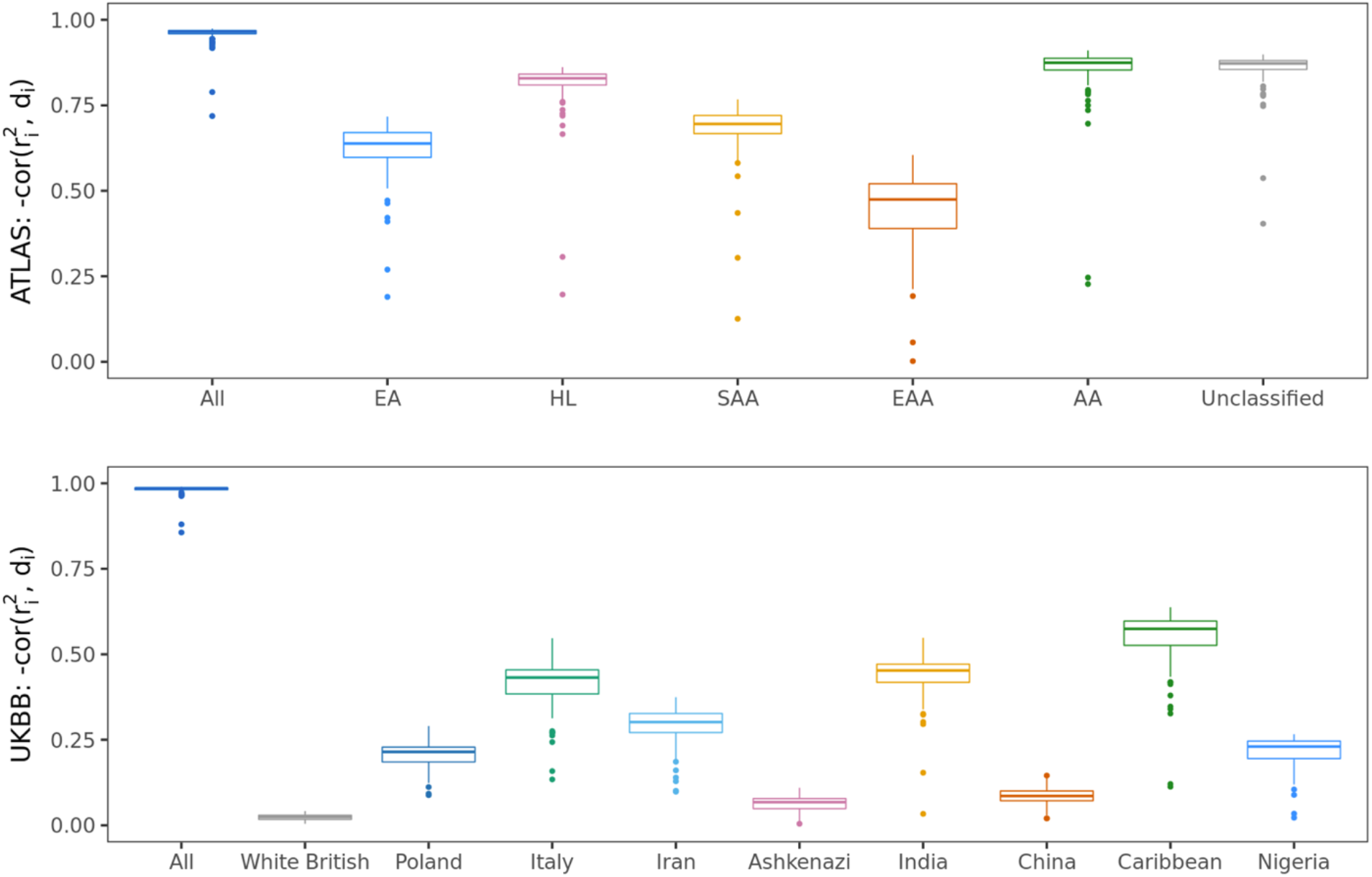
The correlation between individual PGS accuracy and genetic distance is pervasive across 84 traits across ATLAS and UKBB. (a) The distribution of correlation between PGS accuracy and genetic distance for 84 traits in ATLAS. (b) The distribution of correlation between PGS accuracy and genetic distance for 84 traits in UKBB. Each boxplot contains 84 points corresponding to the correlation between PGS accuracy and genetic distance within the group specified by x-axis for each of the 84 traits. The box shows the first, second and third quartile of the 84 correlations, and whiskers extend to the minimum and maximum estimates located within 1.5 × IQR from the first and third quartiles, respectively. Numerical results are reported in Supplementary Table 2 and 3.

### Genetic distance correlates with PGS estimates across most traits

We focused so far on investigating the relationship between GD and PGS accuracy. Next, we turn to evaluating the impact of GD on PGS estimates themselves. We find that GD is significantly correlated with PGS estimates for 82 out of 84 traits in UKBB ranging from R=-0.52 to R= 0.74 (Supplementary Figure 4); this broad range of correlations is in stark contrast with the highly consistent negative correlation of GD and PGS 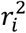. To gain insights into whether PGS coupling with GD is due to stratification or true signal, we next contrasted the correlation of GD to PGS estimates 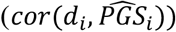 with correlation of GD to the measured phenotype values (*cor*(*d_i_*, y_i_)). We find a wide-range of couplings reflecting trait-specific signals; 30 traits GD correlate in opposite directions with PGS vs. phenotype; 40 trait GD correlates in the same direction with PGS vs. phenotype but differ in correlation magnitude (Supplementary Figure 4). For example, GD shows opposite and significantly different correlations PGS vs trait for years of education (years_of_edu, *cor*(*y_i_,d_i_*) = 0.03, 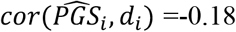). Other traits such as hair color show highly consistent impact of GD on PGS vs trait (darker_hair, *cor*(*y_i_, d_i_*) = 0.59, 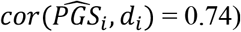; while for monocyte percentage GD shows different magnitudes albeit with the same directions (monocyte_perc, *cor*(*y_i_, d_i_*) =-0.03, 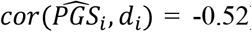). The correlation between GD and phenotype/PGS is also observed in ATLAS. For example, both height phenotype and PGS for height decrease along GD within in ATLAS (Figure 5a); this holds true even if restricted to the European American genetic ancestry cluster (Figure 5b). This is consistent with genetic value driving difference in phenotypes but could also be explained by residual stratification. For neutrophil counts, phenotype and PGS varies in opposite direction along GD across the ATLAS (Figure 5c), although the trend is similar for phenotype and PGS in European American cluster (Figure 5d). This could be explained by genetic value driving signal in Europeans with stratification for other groups. Neutrophil counts have been reported to vary greatly across ancestry groups with reduced counts in individuals of African ancestries^31^. In ATLAS, we observe a negative correlation (−0.04) between GD and neutrophil counts in agreement with the previous reports, while GD is positively correlated (0.08) with PGS estimates with genetically distant individuals traditionally labeled as African American having higher PGS than average. The opposite directions in phenotype/PGS-distance correlations are partly attributed to Duffy-null SNP rs2814778 on chromosome 1q23.2. This variant has a large association with neutrophil counts among individuals traditionally identified as African ancestry, but it is rare and excluded in our training data. This exemplifies the potential bias in PGS due to non-shared causal variants and urges ancestral diversity in genetic studies.

**Figure 5.**
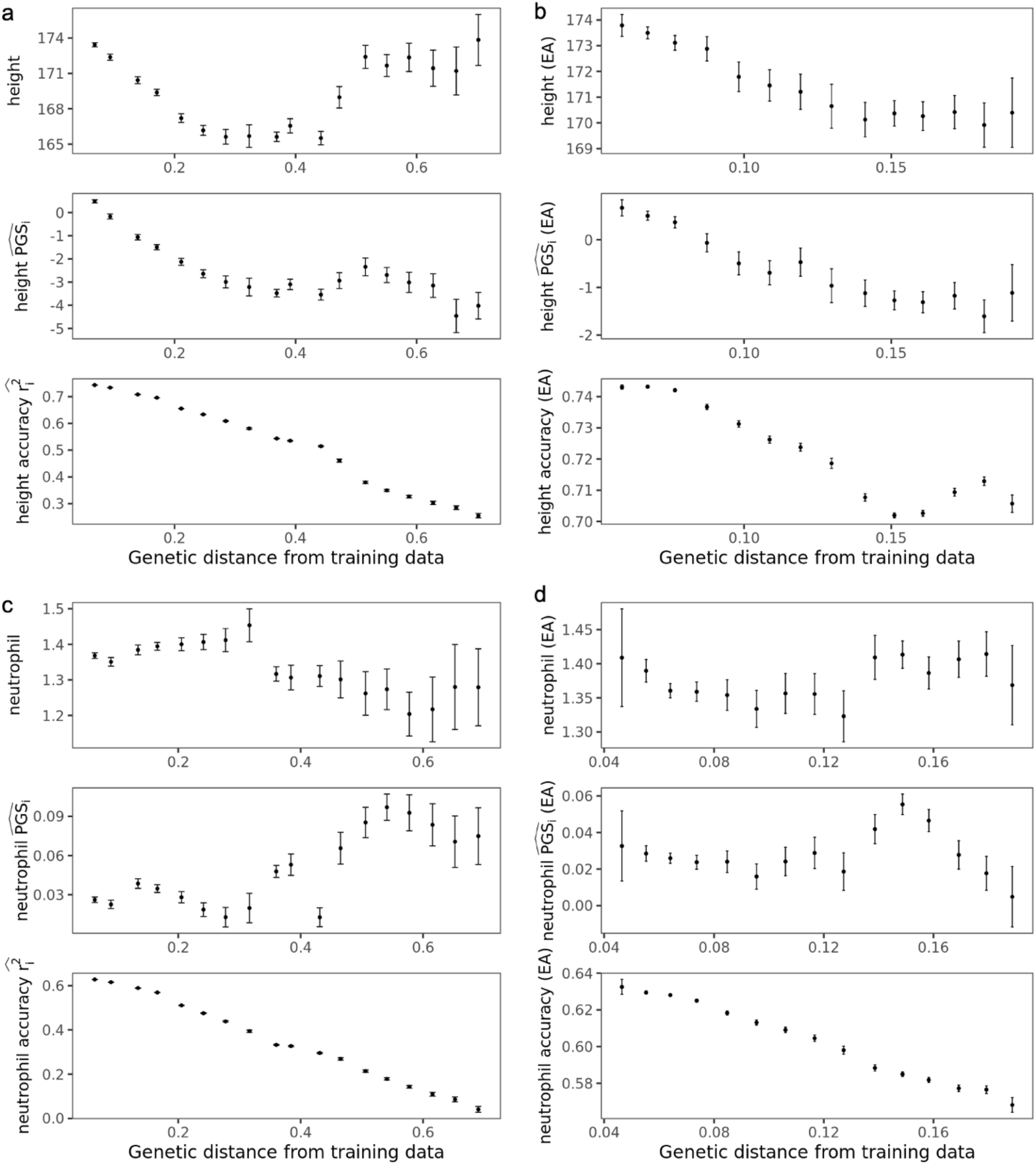
Measured phenotype, PGS estimates, and accuracy varies across the ATLAS and within European American genetic ancestry clusters. (a) Variation of height phenotype, PGS estimates and accuracy across different genetic distance bins in ATLAS. The 36,778 ATALS individuals are divided into 20 equal-interval genetic distance bins. The x-axis is the average genetic distance within the bin, the y-axis is the average phenotype (top), PGS (middle) and individual PGS accuracy (bottom). The error bars represent +/− 1.96 standard error of the mean. Bins with fewer than 50 individuals are not shown due to large standard error of the mean. (b) Variation of height phenotype, PGS estimates and accuracy across different genetic distance bins within European American (EA) genetic ancestry clusters in ATLAS. The 22,380 EA individuals are divided into 20 equal-interval genetic distance bins. The x-axis is the average genetic distance within the bin, the y-axis is the average phenotype (top), PGS (middle) and individual PGS accuracy (bottom). The error bars represent +/− 1.96 standard error of the mean. Bins with fewer than 50 individuals are not shown due to large standard error of the mean. (c) Variation of log neutrophil counts, PGS estimates and accuracy across different genetic distance bins across ATLAS. (d) Variation of log neutrophil counts, PGS estimates and accuracy across different genetic distance bins within European American (EA) genetic ancestry clusters in ATLAS.

Since PGS can vary across GD either as reflection of true signal (i.e. genetic value varying with ancestry) or due to biases in PGS estimation ranging from unaccounted residual population stratification to incomplete data (e.g., partial ancestry-specific tagging of causal effects), our results emphasize the need to consider GD in PGS interpretation beyond adjusting for PGS 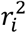.

## Discussion

In this work, we showed that continuous genetic ancestry impacts PGS accuracy and its interpretability across a continuum of genetic ancestries. We proposed individual PGS accuracy as an approach to individualize PGS performance to each target individual. We use a PCA-based genetic distance^23^ from the center of training data to describe an individual’s unique location on the genetic ancestry continuum. In simulations and real data analyses, we showed that individual PGS accuracy tracks well with genetic distance. We demonstrate the pervasive continuous decay of PGS performance as the target individual is further away from the training population. Our demonstration of the continuous PGS accuracy decay directly leads to two conclusions: first, PGS accuracy decay already happens within traditionally defined genetic ancestry groups; second, PGS accuracy can be similar for genetically adjacent individuals that are separated into distinct genetic ancestry groups (usually with different population PGS accuracy). Individual PGS accuracy also enables the evaluation of PGS performance for individuals who cannot be clustered into reference populations, obviating the necessity for genetic ancestry clustering and PGS accuracy evaluation under a discrete view of genetic ancestry.

Our results have several implications for applying PGS to populations with diverse genetic ancestries. First, we highlight the variability in PGS performance along the continuum of genetic ancestry, even within traditionally defined homogenous populations. With the increasing recognition that genetic ancestries are not discrete but rather continuous^15–19^, the individual-level accuracy introduced here provides a powerful tool to study PGS performance along the genetic ancestry continuum. Given the pervasive variable PGS accuracy across individuals, incorporating individual-level metrics of PGS performance can improve the utility of PGS. For example, by using individual-level PGS accuracy, we can identify individuals from Hispanic/Latino genetic ancestry cluster who have similar PGS accuracy with European individuals thus partly alleviating inequities due to lack of access to accurate PGS.

Second, our simulation and real data analysis show that the individual PGS accuracy is highly correlated with genetic distance from training data. The increased genetic distance corresponds to a lower relatedness of the testing individual with the training population^13^. This provides a finer resolution compared with previous theoretical studies that investigate population level PGS portability based on LD/MAF difference, Fst and mean kinship between training and targeting population^12,13^. Along with previous studies, our results emphasize the importance of powerful PGS training in non-European cohorts^32^ to improve the PGS performance for individuals from diverse genetic background. To narrow the prediction gap and ameliorate consequent health disparities between European and non-European ancestries individuals, concerted global effort and equitable collaborations are needed to increase the sample size of underrepresented individuals^32,33^. Equally important is the development of multi-ancestry PGS methods that can effectively leverage ancestrally diverse populations to train PGS models^34^. Some examples of such methods recently developed include PRS-CSx^35^, vilma^36^ and CT-SLEB^37^.

Third, our results highlight the pervasive correlation between PGS estimates and genetic distance from the training data, which usually displays opposite direction or different magnitude compared with the correlation between phenotype and genetic distance. This observation provides a finer resolution of the previously reported mean shift of PGS estimates across genetic ancestry groups^9^. We note that the correlation between genetic distance and phenotype can stem from both potential bias in PGS estimates and true biological difference such as continuous genetic variation. We provide neutrophil counts PGS as an example of potential bias due to low allele frequency of Duffy-null SNP rs2814778^31^ in the training data, however we cannot rule out the impact of true biological differences for most traits. More effort is needed to investigate the PGS bias especially in the context of continuous genetic ancestry.

We note several limitations and future directions of our work. First, individual PGS accuracy is derived from individual PGS uncertainty with approximations under strong assumptions that the causal variants and effects are the same across all genetic ancestries. In reality, despite the abundance of shared causal variants^38^ and the strong transethnic genetic effect sizes correlation^39^, population-specific causal variants and effects still exist and limit the transferability of PGS. Future work could investigate the impact of the population-specific components of genetic architecture on the calibration of PGS accuracy. Second, we approximate the variance of genetic value with heritability and set the value fixed for all individuals. Further work can be done to quantify the genetic value variance for individuals at different genetic distance and assess its impact on accuracy. Third, individual PGS accuracy evaluates how well PGS estimates the genetic value instead of how accurate the PGS predicts the phenotype. Quantifying the individual accuracy of PGS with respect to phenotype can be achieved by modeling environments to calibrate over phenotypes. Fourth, there can be misspecification of model assumptions for the individual-level PGS uncertainty. Future work can be done to investigate the impact of the genetic architectures on the calibration of PGS uncertainty/accuracy. Fifth, limited by the sample size, we train PGS on white British individuals in UKBB, and inevitably define genetic distance relative to European individuals. This work should be replicated while training PGS in non-European individuals in future works. Alternative definitions of genetic distance such as genetic relatedness^13^ and other multi-dimensional descriptions of genetic ancestry continuum^16^ can also be explored in the future.

## Methods

### Model

We model the phenotype of an individual with a standard linear model 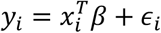 where *x_i_* is an *M* × 1 genotype vector indicating allele counts *β* is an *M* × 1 vector of allelic genetic effects and *ϵ_i_* is random noise. Under a random effects model, *β* is a vector of random variable sampled from a prior distribution *p*(*β*) which differs under different genetic architecture assumptions^40^ and PGS methods^29,41–43^. The PGS weights 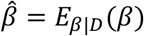 are estimated to be the posterior mean given the observed data *D* (*D* = (*X_train_, y_train_*) with access to individual-level genotypes *X_train_* and phenotype *y_train_* or 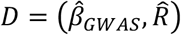 with access to marginal association statistics 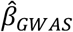 and 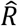). An individual i’s genetic value 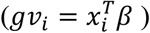 is estimated to be 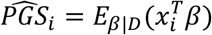, the uncertainty of which is estimated as the posterior variance of genetic value 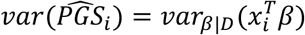^26^.

### Individual PGS accuracy

Under a random effects model both the genetic value and PGS estimate for individual *i*. are random variables. The randomness of 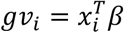 comes from the randomness in *β* and the randomness of 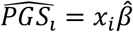 comes from the randomness of both *β* and the training data *D*. Individual PGS accuracy measures the correlation between *gv_i_* and 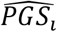, which can be computed with the following equation:

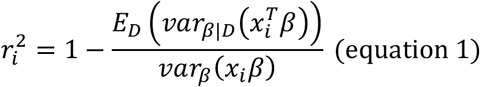

where 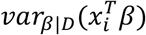 is the posterior variance of genetic value given the training data and 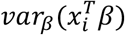 is the genetic variance. The equation is derived as follows:

First, we show that under the random effects model, 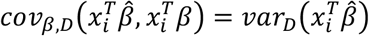 (where 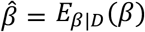) following equation 5.149 in ref^44^:

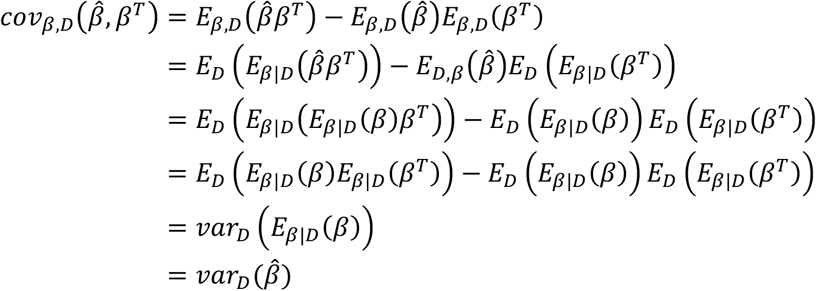

Multiply *x_i_* on both sides of equation, we obtain:

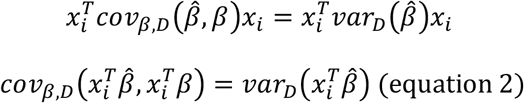

Next, by applying the law of total variance, we show that:

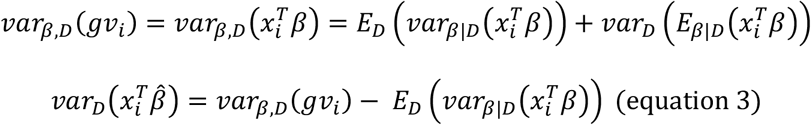

Third, we derive the correlation between *gv_i_* and 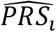 as:

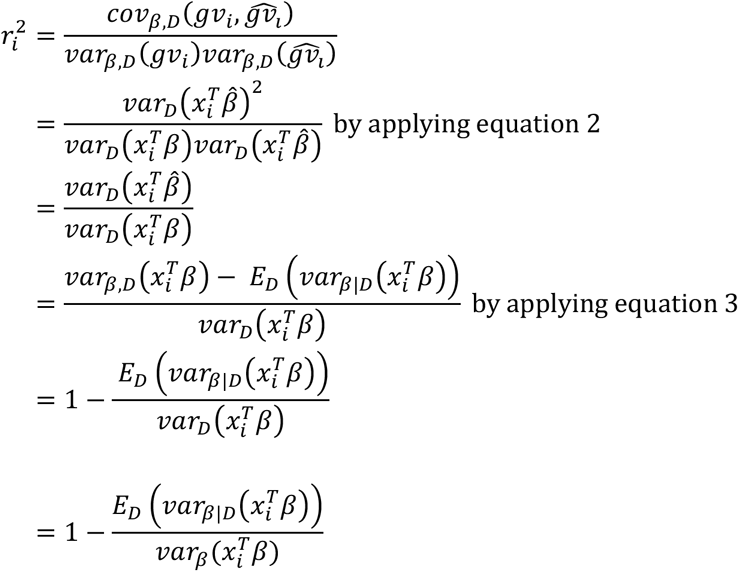

Equation 1 is widely used in animal breeding theory to compute the reliability of estimated breeding value for each individual^30^. In this work, we use individual PGS uncertainty 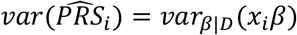 as an unbiased estimator of 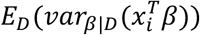. We also use estimated heritability to approximate 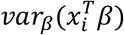 in simulation where the phenotype has unit variance. In real data analysis, since the phenotypes does not necessarily have unit variance, we approximate 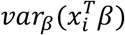 by scaling the estimated heritability multiplied by the residual phenotypic variance in the training population after regressing GWAS covariates including sex, age and precomputed UKBB PC1-16 (Data-Field 22009).

### Analytical form of individual PGS accuracy under infinitesimal assumption

Without loss of generality, we assume a prior distribution of genetic effects as follows:

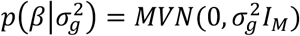

With access to individual genotype data *X_train_* and phenotypes *y_train_*, the likelihood of the data is

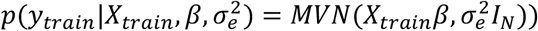

The posterior distribution of genetic effects given the data is proportional to the product of the prior and the likelihood:

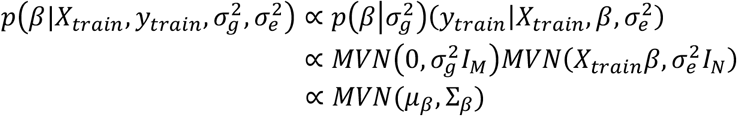

Where 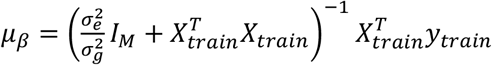 and 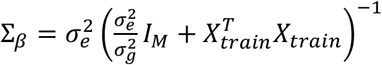

For a new target individual, the posterior variance of the genetic value is:

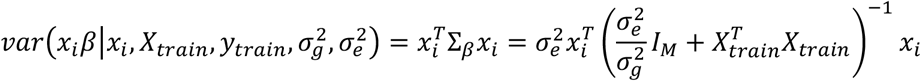

After performing eigendecomposition on 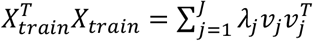, we can rewrite

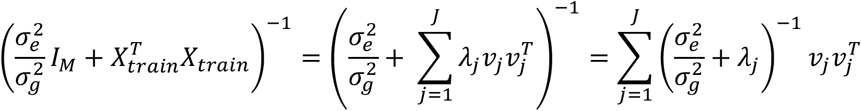

where *v_j_* and *λ_j_* corresponds to the *j_th_* eigenvalue and unit-length eigenvector of training genotype *X_train_*

Thus, we can rewrite the posterior variance of genetic value as

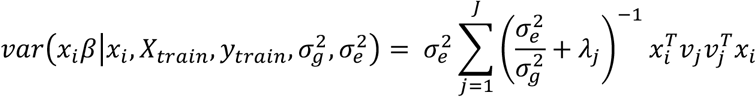

Replacing 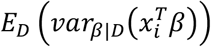 in equation 1 with analytical form of 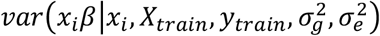, we get

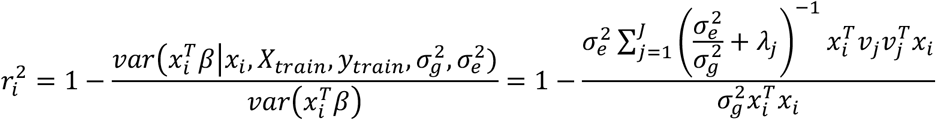

As the eigenvalue of 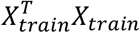 increases linearly with training sample size N^45^. At the UKBB level sample size (e.g. N = 371,018for our UKBB white British training data), the eigenvalue for the top PCs are usually larger than the ratio of environmental noise variance and genetic variance 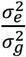. Thus, we can further approximate the analytical form with:

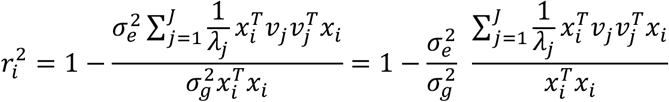

The term 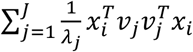 is the Mahalanobis distance of the testing individual *í* from the center of the training genotype data on its PC space and 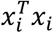 is the sum of squared allelic counts across all variants.

Empirically, the ratio between the two is highly correlated with the Euclidean distance of the individual from the training data on that PC space (R= 1, P-value < 2.2e-16 in UKBB).

### Genetic Distance

The genetic distance is defined as the Euclidean distance between a target individual and the center of training data on the PC space of training data.

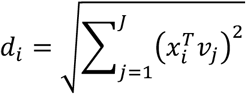

where *d_i_* is the genetic distance of a testing individual *i*. from the training data, *x_i_* is an M x1 genotype vector for testing individual *i*, *v_j_* is the *j_th_* eigenvector for the genotype matrix of training individuals and *J* is set to 20.

### Ancestry ascertainment in UKBB

The UKBB individuals are clustered into nine sub-continental ancestry clusters White British, Poland, Iran, Italy, Ashkenazi, India, China, Caribbean and Nigeria based on the top 16 precomputed PCs (Data-Field 22009) as described in ref^23^. The center of each ancestry group on PC space is obtained from ref^23^. Each individual is assigned to one of the nine ancestral groups based on their Euclidean distance to the centers on the PC space. The genetic ancestry of an individual is labeled as unknown if its distance to any genetic ancestry center is larger than one eighth of maximum distance between any pairs of cub-continental ancestry clusters. We are able to cluster 91% ofthe UKBB participants into 411,018 British white, 4127 Polish, 1169 Iran, 6499 Italy, 2352 Ashkenazi, 1798 China, 2472 Caribbean and 3894 Nigeria.

### Ancestry ascertainment in ATLAS

The ATLAS individuals are clustered into five genetic ancestry clusters - European Americans (EA), Hispanic and Latino Americans (HL), South Asian Americans (SAA) and East Asian Americans (ESA) and African Americans (AA) as described in ref^25^ based on their proximity with 1000 Genome super populations on the PC space. First, we filter the ATLAS typed genotypes with plink2 by Mendel error rate (‘plink --me 1 1 –set-me-missing’), founders (‘--filter-founders’), minor allele frequency (‘–maf 0.15’), genotype missing call rate (‘--geno 0.05’), and Hardy-Weinberg equilibrium test p-value (‘-hwe 0.001’). Next, ATLAS genotypes were merged with the 1000 Genomes phase 3 dataset. Then, LD pruning was performed on the merged dataset (‘--indep 200 5 1.15 -- indep-pairwise 100 5 0.1’). The top10 PCs were computed with the flashpca2^46^ software with all default parameters. Next, we use the super population label and PCs of the 1000 Genome individuals to train the K-nearest neighbors (KNN) model to assign genetic ancestry labels to each ATLAS individual. For each ancestry cluster, we run KNN on the pair of PCs that capture the most variation for each genetic ancestry group: European, East Asian, and African ancestry groups utilize PCs 1 and 2, the Admixed American group use PCs 2 and 3, and the South Asian group use PCs 4 and 5. In each analysis, we use 10-fold cross-validation to select the ‘k’ hyper-parameter from k=5, 10, 15, 20. If an individual is assigned to multiple ancestries with probability larger than 0.5 or is not assigned to any clusters, it’s labeled as unknown. We relabel the five 1000 genome super population as EA for EUR, HL for AMR, SAA for SAS, AA for AFR and ESA for EAS. We can cluster 95% of the ATLAS participants into 22,380 EA, 6973 HL, 625 SAA, 3331 EAA, 1995 AA and 2332 individuals are labeled as unknown.

### Genotype data

In simulations, we use 1,054,151 UKBB HapMap3 SNPs for simulating phenotypes, training PGS models and calculating PGS for testing individuals in UKBB. For real data analysis, we use an intersection of UKBB HapMap 3 SNPs and ATLAS imputed SNPs for the training of PGS in UKBB and calculating PGS for remaining UKBB individuals and ATLAS individuals. We start from 1,054,151 UKBB HapMap3 SNPs and 8,048,268 ATLAS imputed SNPs. Since UKBB is on genome build hg37 and ATLAS is on hg38, we first lift all ATLAS SNPs from hg38 to hg37 with *snp_modifyBuild* function in bigsnpr R package. Next, we match UKBB SNPs and ATLAS SNPs by chromosome and position with *snp_match* function in bigsnpr. Then, we recode ATLAS SNPs using UKBB reference alleles with plink2 --recode flag. In the end, 979,457 SNPs remain for training the LDpred2 models in real data analysis.

### Simulated phenotypes

We use simulations on all UKBB individuals to investigate the impact of genetic distance from training data on the various metrics of PGS. We fix the proportion of causal SNPs *p_causal_* = 0.01 and heritability as 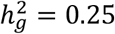. The simulated genetic effects and phenotype are generated as follows: First, we randomly sample

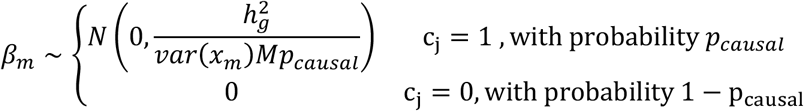

where *var*(*x_m_*) is the variance of allele counts for SNP *m* among all UKBB individuals. Second, we compute the genetic value for each individual as 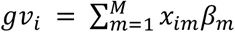 and randomly sample environmental noise 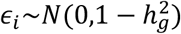. Third, generate phenotype as *y_i_* = *gv_i_* + *ϵ_i_*. We repeat the process 100 times to generate 100 sets of genetic values and phenotypes.

### Genetic distance from PGS training data

To compute the genetic distance of testing individuals from the training population, we perform PCA on the 371,018 UKBB white British training individuals and project the 48,586 UKBB testing individuals and 36,778 ATLAS training individuals on the PC space. We start from the 979,457 SNPs that are overlapped in UKBB and ATLAS. First, we perform LD pruning with plink2 (--indep-pairwise 1000 50 0.05) and exclude the long-range LD regions. Next, we perform PCA analysis with flashpca2^46^ on the 371,018 UKBB white British training individuals to obtain the top 20 PCs. Then, we project the remaining 48,586 UKBB individuals that are not included in the training data and 36,778 ATLAS individuals onto the PC space of training data by using SNP loadings (--outload loadings.txt) and their means and standard deviations (--outmeansd meansd.txt) output from flashpca2. In the end, we compute the genetic distance for each individual as the norm of its projection on the PC space.

### LDpred2 PGS model training

The PGS models were trained on 371,018 UKBB individuals labeled as white British with the LDpred2^29^ method for both simulation and real data analysis. For simulation analysis, we use 1,054,151 UKBB HapMap3 variants. For real data analysis, we use 979,457 SNPs that are overlapped in UKBB HapMap3 variants and ATLAS imputed genotypes.

First, we obtain GWAS summary statistics by performing GWAS on the training individuals with plink2 using sex, age and precomputed PC-1-16 as covariates. Second, we calculate the in-sample LD matrix with the function snp_cor from R package bigsnpr^47^. Next, we use the GWAS summary statistics and LD matrix as input for *snp_ldpred2_auto* function in bigsnpr to sample from the posterior distribution of genetic effect sizes. Instead of using a held-out validation dataset to select hyperparameters p (proportion of causal variants) and h2 (heritability), *snp_ldpred2_auto* estimates the two parameters from data with MCMC directly. We run 10 chains with different initial sparsity p from 10^-4^ to 1 equally spaced in log space. For all chains, we set the initial heritability as the LD score regression heritability^48^ estimated by the built-in function *snp_ldsc.* We perform quality control of the 10 chains by filtering out trains with estimated heritability that are smaller than 0.7 times of the median heritability of the 10 chains or with estimated sparsity that are smaller than 0.5 times of the median sparsity or 2 times of the median sparsity. For each chain that passes filtering, we remove the first 100 MCMC iterations as burn-in and thin the next 500 iterations by selecting every 5th iteration to reduce autocorrelation between MCMC samples. In the end, we obtain a M x B matrix 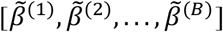, where each column of the matrix 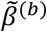 is a sample of posterior causal effects of the M SNPs. Due to the quality control of MCMC chains, the total number of posterior samples *B* ranges from 500 to 1000.

### Calculate PGS and accuracy

We use the score function in plink2 to compute the PGS for 48,586 and 36,778 testing individuals in UKBB and ATLAS, respectively. For each 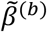 we compute the PGS for each individual i as 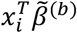 with plink2 (--score). For each individual with genotype *x_i_*, we compute 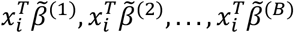 to approximate its posterior distribution of genetic value. The genotype 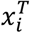 is centered to the average allele count (--read-freq) in training data to reduce the uncertainty from the unmodeled intercept. We estimate the PGS with the posterior mean of the genetic value as 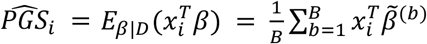. We estimate the individual level PGS uncertainty as 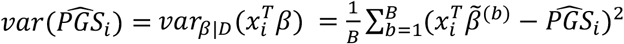. The individual level PGS accuracy is calculated as 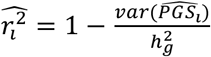 for simulation (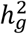 is the heritability estimated by the LDpred2 model) and as 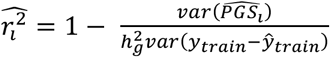 for real data analysis, where *var*(*y_train_* – *ŷ_train_*) refers to the variance of residual phenotype in training data after regressing out GWAS covariates).

### Calibration of credible interval in simulation

We run the LDpred2 model on 371,018 white British training individuals for the 100 simulation replicates. In each simulation *r*, for individual with genotype *x_i_*, we compute 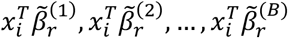 to approximate its posterior distribution of genetic value, generate 90% credible interval *CI* – *GV_ir_* with 5% and 95% quantile of the distribution and check if its genetic value is contained in the credible interval *I*(*gv_ir_* ∈ *CI* – *GV_ir_*). We compute the empirical coverage for each individual as the mean across the 100 simulation replicates 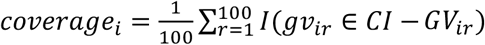.

## Ethics declarations

All research performed in this study conformed with the principles of the Helsinki Declaration. All individuals provided written informed consent to participate in the study. Patient Recruitment and Sample Collection for Precision Health Activities at UCLA is an approved study by the UCLA Institutional Review Board (UCLA IRB). IRB#17-001013.

## Data availability

The individual-level genotype and phenotype data are available by application from the UKBB http://www.ukbiobank.ac.uk/. Summary statistics for UCLA ATLAS data are available at https://atlas-phewas.mednet.ucla.edu/.

## URLs

LDPred2 software implementing individual PRS uncertainty: https://privefl.github.io/bigsnpr/articles/LDpred2.html

## Acknowledgments

This research was conducted using the UK Biobank Resource under application 33297. We thank the participants of UK Biobank for making this work possible. This work was funded in part by NIH awards U01HG011715, R01HG009120, R01MH115676.The content is solely the responsibility of the authors and does not necessarily represent the official views of the NIH.

## Supplementary Figures

**Supplementary Figure 1.**
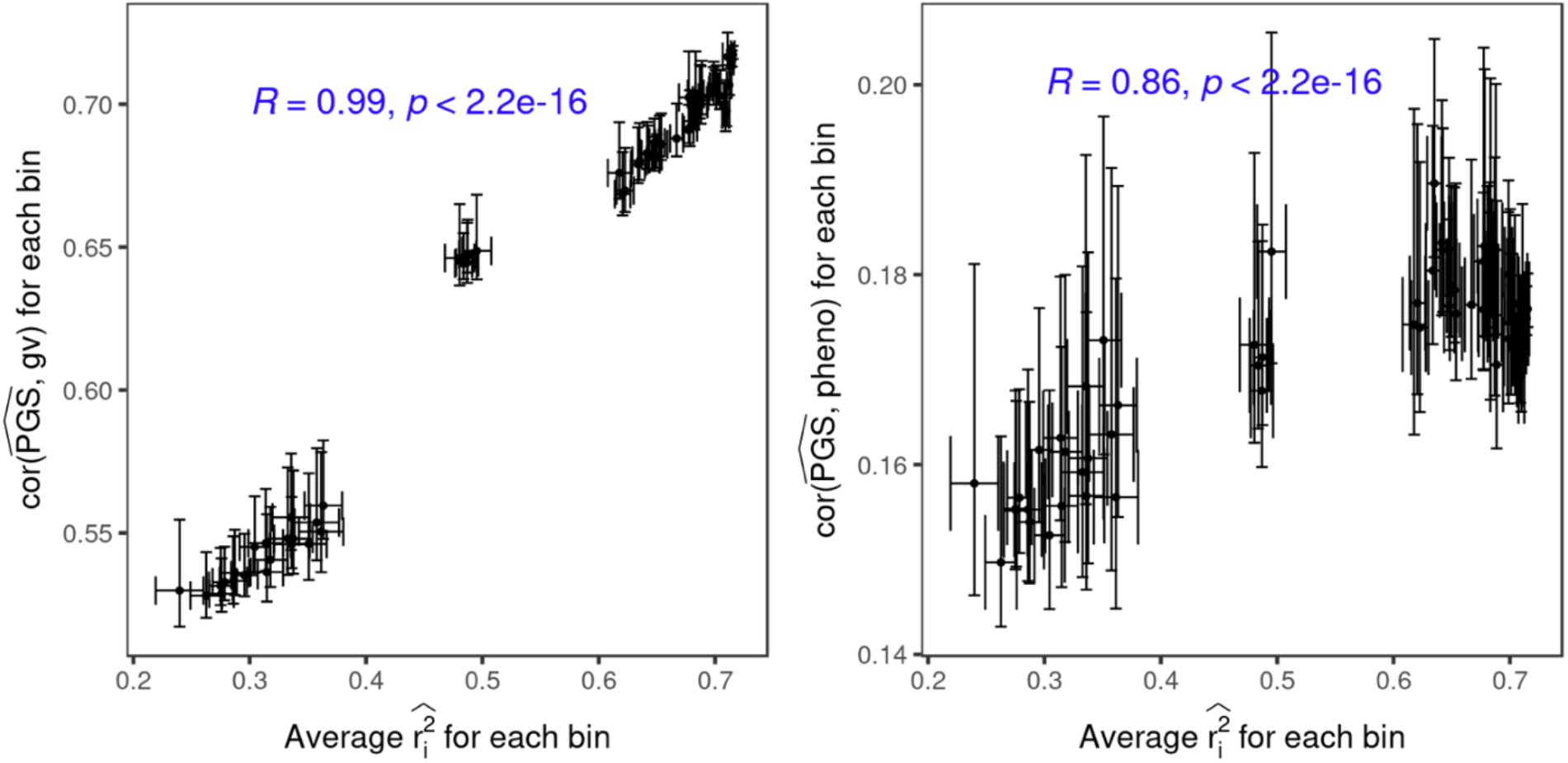
The individual level accuracy is highly correlated with population level accuracy. All UKBB testing individuals are divided into 100 bins based on their genetic distance. The x-axis is the average individual-level PGS accuracy for the individuals within the bin and the y-axis is (a) the squared correlation between simulated genetic value and PGS estimates for the individuals within the bin (b) the squared correlation between simulated phenotype and PGS estimates. The dot and error bars represent the mean and ± 1.96 s.e.m from 100 simulations.

**Supplementary Figure 2.**
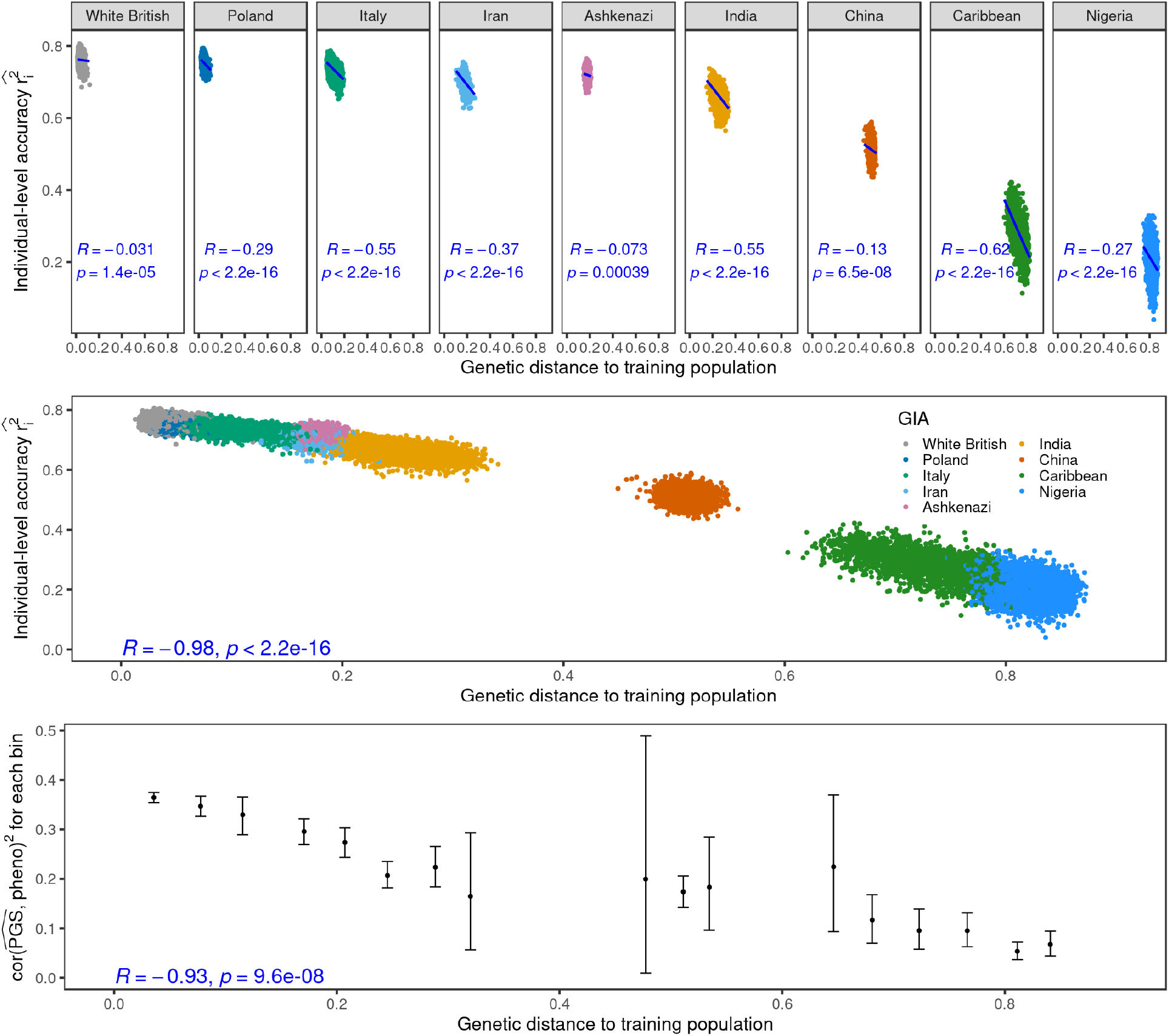
The individual-level accuracy for height PGS decreases across the genetic ancestry continuum in UKBB. (a) Individual PGS accuracy decreases within subcontinental admixed genetic ancestry clusters. Each dot represents a testing individual from UKBB. For each dot, the x-axis represents its distance from the training population on the genetic continuum; the y-axis represents its PGS accuracy. The color represents the inferred genetic ancestry cluster. R and p refer to the correlation between genetic distance and individual-level PGS accuracy and its significance from two-sided t-tests. (b) Individual PGS accuracy decreases across the entire UKBB. (c) The population PGS accuracy decreases with the average genetic distance in each genetic distance bin. All UKBB individuals are divided into 20 equal-interval genetic distance bins. The x-axis is the average genetic distance within the bin; the y-axis is the squared correlation between PGS and phenotype for individuals in the bin. The dot and error bar show mean and 95% confidence interval from 1000 bootstrap samples.

**Supplementary Figure 3.**
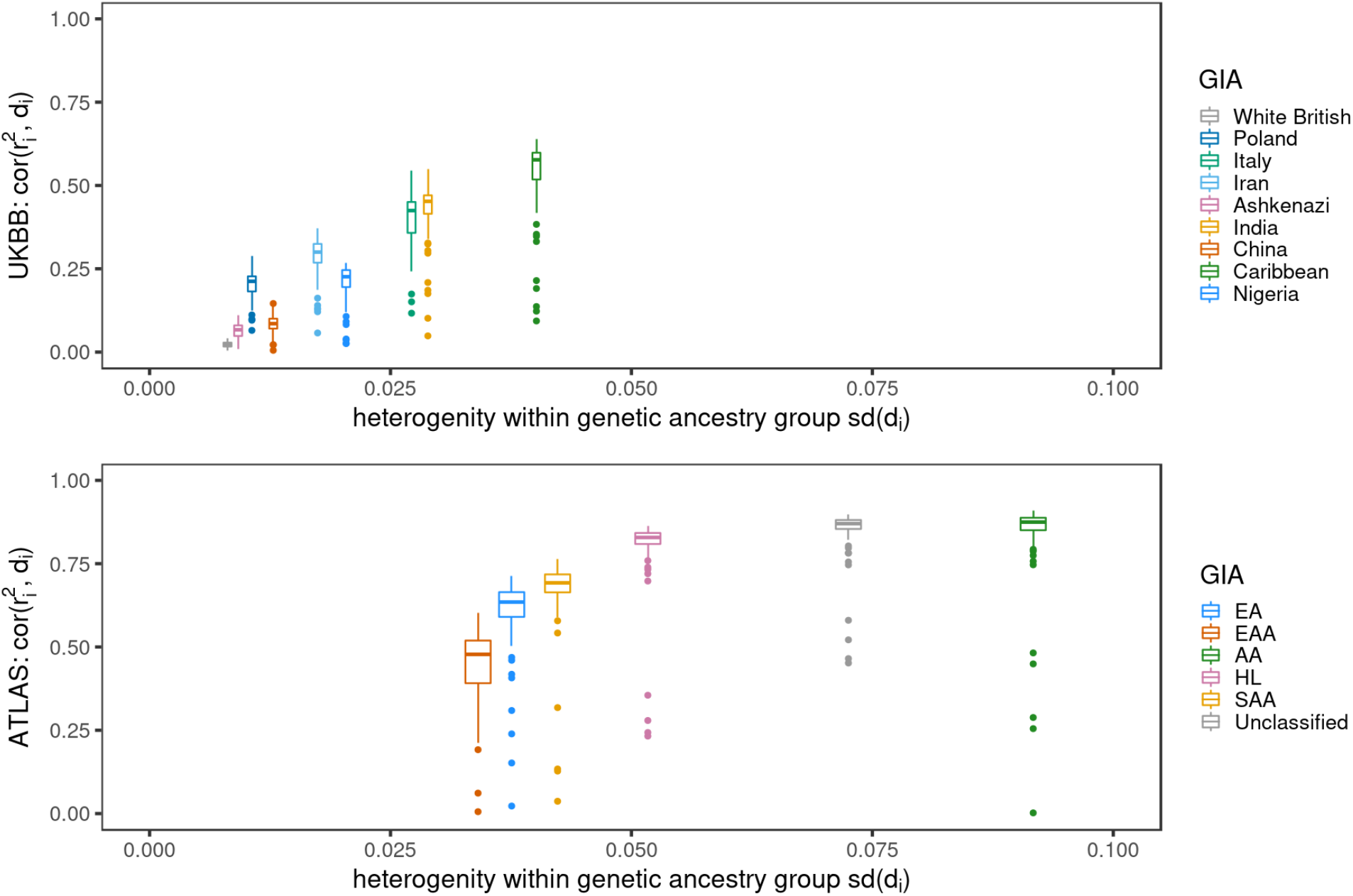
Lower heterogeneity within the genetic ancestry group corresponds to a lower correlation between genetic distance and individual PGS accuracy. (a) The distribution of correlations between PGS accuracy and genetic distance for 84 traits in ATLAS. (b)The distribution of correlations between PGS accuracy and genetic distance for 84 traits in UKBB. The x-axis is the homogeneity of the genetic ancestry clusters measured as standard deviation of genetic distance within a genetic ancestry cluster; a larger *sd*(*d_i_*) indicates a larger variation of genetic background. Each boxplot contains 84 points corresponding to the correlation between PGS accuracy and genetic distance within the group specified by x-axis for each of the 84 traits. The box shows the first, second and third quartile of the 84 correlations, and whiskers extend to the minimum and maximum estimates located within 1.5 × IQR from the first and third quartiles, respectively.

**Supplementary Figure 4.**
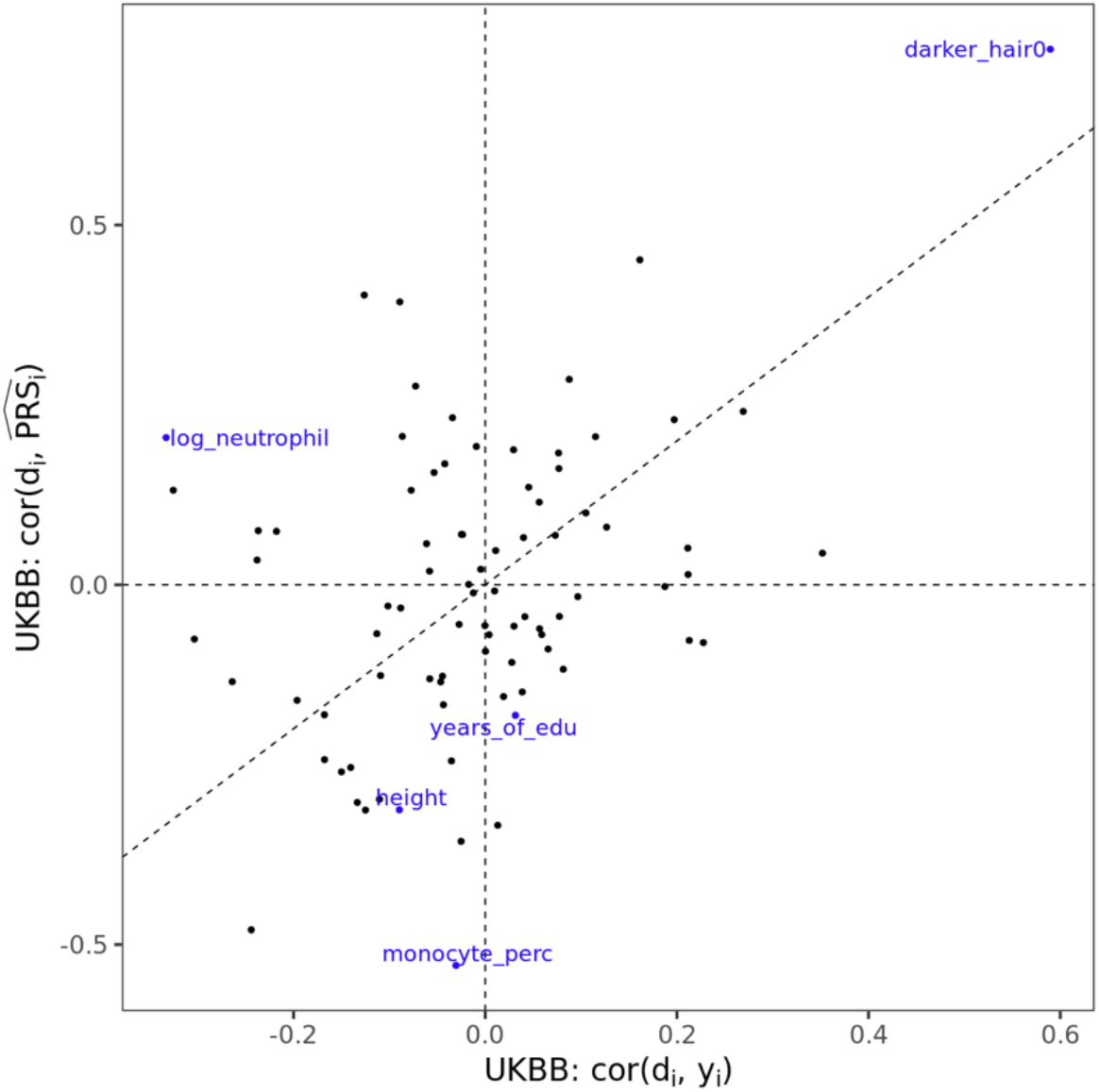
Discordant directions of phenotype/PGS-distance correlations in UKBB. The x axis is the correlation between phenotype and genetic distance and the y axis is the correlation between PGS estimates and genetic distance for all 48,586 testing individuals in UKBB. Numerical results are reported in Supplementary Table 4.

**Supplementary Table 1. The training sample size, proportion of causal variants and heritability of the 84 traits.**

**Supplementary Table 2. The correlation between individual PGS accuracy and genetic distance from training data across ATLAS and within each genetic ancestry clusters**

**Supplementary Table 3. The correlation between individual PGS accuracy and genetic distance from training data across UKBB and within each genetic ancestry clusters**

**Supplementary Table 4. The correlation between measured phenotype/PGS and genetic distance from training data across UKBB**

## Reference

1. Torkamani, A., Wineinger, N. E. & Topol, E. J. The personal and clinical utility of polygenic risk scores. Nat. Rev. Genet. 19, 581–590 (2018).

2. Lambert, S. A., Abraham, G. & Inouye, M. Towards clinical utility of polygenic risk scores. Hum. Mol. Genet. 28, R133–R142 (2019).

3. Kullo, I. J. et al. Polygenic scores in biomedical research. Nat. Rev. Genet. (2022) doi:10.1038/s41576-022-00470-z.

4. Li, R., Chen, Y., Ritchie, M. D. & Moore, J. H. Electronic health records and polygenic risk scores for predicting disease risk. Nat. Rev. Genet. 21, 493–502 (2020).

5. Natarajan, P. et al. Polygenic Risk Score Identifies Subgroup With Higher Burden of Atherosclerosis and Greater Relative Benefit From Statin Therapy in the Primary Prevention Setting. Circulation 135, 2091–2101 (2017).

6. Lee, A. et al. BOADICEA: a comprehensive breast cancer risk prediction modelincorporating genetic and nongenetic risk factors. Genet. Med. 21, 1708–1718 (2019).

7. Khera, A. V. et al. Polygenic Prediction of Weight and Obesity Trajectories from Birth to Adulthood. Cell 177, 587–596.e9 (2019).

8. Perkins, D. O. et al. Polygenic Risk Score Contribution to Psychosis Prediction in a Target Population of Persons at Clinical High Risk. Am. J. Psychiatry 177, 155–163 (2020).

9. Martin, A. R. et al. Human Demographic History Impacts Genetic Risk Prediction across Diverse Populations. Am. J. Hum. Genet. 100, 635–649 (2017).

10. Martin, A. R. et al. Clinical use of current polygenic risk scores may exacerbate health disparities. Nat. Genet. 51, 584–591 (2019).

11. Mostafavi, H. et al. Variable prediction accuracy of polygenic scores within an ancestry group. Elife 9, e48376 (2020).

12. Wang, Y. et al. Theoretical and empirical quantification of the accuracy of polygenic scores in ancestry divergent populations. Nat. Commun. 11, 3865 (2020).

13. Scutari, M., Mackay, I. & Balding, D. Using Genetic Distance to Infer the Accuracy of Genomic Prediction. PLoS Genet. 12, e1006288 (2016).

14. Duncan, L. et al. Analysis of polygenic risk score usage and performance in diverse human populations. Nat. Commun. 10, 1–9 (2019).

15. Coop, G. Genetic similarity and genetic ancestry groups. arXiv [q-bio.PE] (2022).

16. Lewis, A. C. F. et al. Getting genetic ancestry right for science and society. Science 376, 250–252 (2022).

17. Mathieson, I. & Scally, A. What is ancestry? PLoS Genet. 16, e1008624 (2020).

18. Krainc, T. & Fuentes, A. Genetic ancestry in precision medicine is reshaping the race debate. Proceedings of the National Academy of Sciences 119, e2203033119 (2022).

19. Belbin, G. M. et al. Toward a fine-scale population health monitoring system. Cell 184, 2068–2083.e11 (2021).

20. Wojcik, G. L. et al. Genetic analyses of diverse populations improves discovery for complex traits. Nature 570, 514–518 (2019).

21. Bitarello, B. D. & Mathieson, I. Polygenic Scores for Height in Admixed Populations. G3 10, 4027–4036 (2020).

22. Clarke, S. L. et al. Race and Ethnicity Stratification for Polygenic Risk Score Analyses May Mask Disparities in Hispanics. Circulation 146, 265–267 (2022).

23. Privé, F. et al. Portability of 245 polygenic scores when derived from the UK Biobank and applied to 9 ancestry groups from the same cohort. Am. J. Hum. Genet. 109, 12–23 (2022).

24. Johnson, R. D. et al. The UCLA ATLAS Community Health Initiative: promoting precision health research in a diverse biobank. medRxiv (2022).

25. Johnson, R. et al. Leveraging genomic diversity for discovery in an electronic health record linked biobank: the UCLA ATLAS Community Health Initiative. Genome Med. 14, 1–23 (2022).

26. Ding, Y. et al. Large uncertainty in individual polygenic risk score estimation impacts PRS-based risk stratification. Nat. Genet. 54, 30–39 (2021).

27. Price, A. L. et al. Principal components analysis corrects for stratification in genome-wide association studies. Nat. Genet. 38, 904–909 (2006).

28. Alexander, D. H. & Lange, K. Enhancements to the ADMIXTURE algorithm for individual ancestry estimation. BMC Bioinformatics 12, 246 (2011).

29. Privé, F., Arbel, J. & Vilhjálmsson, B. J. LDpred2: better, faster, stronger. Cold Spring Harbor Laboratory 2020.04.28.066720 (2020) doi:10.1101/2020.04.28.066720.

30. Walsh, B. & Lynch, M. Evolution and Selection of Quantitative Traits. in Evolution and Selection of Quantitative Traits (Oxford University Press, 2018).

31. Reich, D. et al. Reduced neutrophil count in people of African descent is due to a regulatory variant in the Duffy antigen receptor for chemokines gene. PLoS Genet. 5, e1000360 (2009).

32. Fatumo, S. et al. A roadmap to increase diversity in genomic studies. Nat. Med. 28, 243–250 (2022).

33. Martin, A. R. et al. Increasing diversity in genomics requires investment in equitable partnerships and capacity building. Nat. Genet. 54, 740–745 (2022).

34. Wang, Y., Tsuo, K., Kanai, M., Neale, B. M. & Martin, A. R. Challenges and Opportunities for Developing More Generalizable Polygenic Risk Scores. Annu. Rev. Biomed. Data Sci. 5, 293–320 (2022).

35. Ruan, Y. et al. Improving polygenic prediction in ancestrally diverse populations. Nat. Genet. 54, 573–580 (2022).

36. Spence, J. P., Sinnott-Armstrong, N., Assimes, T. L. & Pritchard, J. K. A flexible modeling and inference framework for estimating variant effect sizes from GWAS summary statistics. bioRxiv 2022.04.18.488696 (2022) doi:10.1101/2022.04.18.488696.

37. Zhang, H. et al. Novel Methods for Multi-ancestry Polygenic Prediction and their Evaluations in 3.7 Million Individuals of Diverse Ancestry. bioRxiv 2022.03.24.485519 (2022) doi:10.1101/2022.03.24.485519.

38. Shi, H. et al. Localizing Components of Shared Transethnic Genetic Architecture of Complex Traits from GWAS Summary Data. Am. J. Hum. Genet. 106, 805–817 (2020).

39. Shi, H. et al. Population-specific causal disease effect sizes in functionally important regions impacted by selection. Nat. Commun. 12, 1098 (2021).

40. Hou, K. et al. Accurate estimation of SNP-heritability from biobank-scale data irrespective of genetic architecture. Nat. Genet. 51, 1244–1251 (2019).

41. Ge, T., Chen, C.-Y., Ni, Y., Feng, Y.-C. A. & Smoller, J. W. Polygenic prediction via Bayesian regression and continuous shrinkage priors. Nat. Commun. 10, 1776 (2019).

42. Zhang, Q., Privé, F., Vilhjálmsson, B. & Speed, D. Improved genetic prediction of complex traits from individual-level data or summary statistics. Cold Spring Harbor Laboratory 2020.08.24.265280 (2020) doi:10.1101/2020.08.24.265280.

43. Lloyd-Jones, L. R. et al. Improved polygenic prediction by Bayesian multiple regression on summary statistics. Nat. Commun. 10, 1–11 (2019).

44. Sorensen, D. & Gianola, D. Likelihood, Bayesian, and MCMC Methods in Quantitative Genetics. (Springer New York).

45. Abraham, G. & Inouye, M. Fast principal component analysis of large-scale genome-wide data. PLoS One 9, e93766 (2014).

46. Abraham, G., Qiu, Y. & Inouye, M. FlashPCA2: principal component analysis of Biobank-scale genotype datasets. Bioinformatics 33, 2776–2778 (2017).

47. Privé, F., Aschard, H., Ziyatdinov, A. & Blum, M. G. B. Efficient analysis of large-scale genomewide data with two R packages: bigstatsr and bigsnpr. Bioinformatics 34, 2781–2787 (2018).

48. Bulik-Sullivan, B. K. et al. LD Score regression distinguishes confounding from polygenicity in genome-wide association studies. Nat. Genet. 47, 291–295 (2015).

